# Great tits differ in glucocorticoid plasticity in response to spring temperature

**DOI:** 10.1101/2022.04.21.489013

**Authors:** Michaela Hau, Caroline Deimel, Maria Moiron

## Abstract

Fluctuations in environmental temperature affect energy metabolism, stimulating the expression of phenotypic plasticity in behavioral and physiological traits. Vertebrate hormonal signals like glucocorticoids underpin environmentally-induced phenotypic plasticity, with changes in circulating concentrations orchestrating plastic changes in diverse traits. Climate change is predicted to alter temperature variation globally, making it imperative to evaluate whether free-living animal populations can cope adaptively. To evaluate their potential to respond to ongoing global change, we quantified individual variation in glucocorticoid plasticity to ambient temperature in wild great tits (*Parus major*). Using a reaction norm approach, we repeatedly sampled individuals for circulating glucocorticoid concentrations across five years. As expected, baseline and stress-induced glucocorticoid concentrations increased with lower ambient temperatures at the population and within-individual level. Moreover, we provide unique evidence that free-living individuals differ significantly in their plastic responses to temperature variation for both glucocorticoid traits, with some displaying greater plasticity than others. Average concentrations and degree of plasticity covaried for baseline glucocorticoids, indicating that these two reaction norm components are linked. Hence, individual variation in glucocorticoid plasticity to an important environmental factor exists in a wild population, representing a crucial step to assess the adaptive potential of vertebrates to endure current temperature fluctuations.

## Introduction

Fluctuations in key climatic variables like ambient temperature can affect the energy balance of vertebrate populations (Scholander et al., 1950). To cope with energy-demanding changes in environmental conditions, animals plastically adjust the expression of physiological and behavioral traits. This so-called phenotypic plasticity is a widespread mechanism by which organisms can reversibly and repeatedly respond to fluctuating environments. In response to changes in ambient temperature vertebrates display phenotypic plasticity in an array of thermoregulatory, metabolic and behavioral traits (Angilletta et al., 2010; Kingsolver & Huey, 1998; Tattersall et al., 2012). Such concerted phenotypic plasticity is often orchestrated by hormones, potent internal signals that pleiotropically promote plasticity in a suite of traits (Cox et al., 2016; Ketterson et al., 2009; Wingfield et al., 1998). Glucocorticoids, a conserved set of vertebrate steroid hormones, are particularly sensitive to environmental variation and can undergo rapid changes in circulating concentrations, for example when ambient temperatures fluctuate (Boonstra et al., 2014; de Bruijn & Romero, 2018; Jessop et al., 2016; Wingfield & Ramenofsky, 2011). Glucocorticoid receptors are widely distributed in brain and body tissues, enabling these hormones to function as higher-order regulators (‘integrators’) to pleiotropically promote phenotypic plasticity (Cohen et al., 2012; Martin & Cohen, 2014).

Not all individuals within a single population respond to environmental variation in a similar way. Individuals can differ in how strongly they respond to variation in ambient temperature, for instance in metabolic rate (Briga & Verhulst, 2017; Careau et al., 2014; Kar et al., 2021), foraging activity (Hall & Chalfoun, 2019), or incubation behavior (Cones et al., 2021). Since glucocorticoids support the regulation of such traits (Romero & Wingfield, 2016), quantifying the extent of glucocorticoid plasticity within and among individuals of wild vertebrate populations is crucial for evaluating the potential of these mechanisms to evolve (e.g., Angelier & Wingfield, 2013; Guindre-Parker, 2020; Hau et al., 2016; Malkoc, Mentesana, et al., 2021; Taff & Vitousek, 2016; Wada & Sewall, 2014). Individual differences are the raw material of selection, and for evolution to occur these individual differences have to harbor a heritable component. Indeed, circulating glucocorticoid concentrations are heritable in several vertebrate taxa (e.g., mammals: Bairos-Novak et al., 2018, birds: Béziers et al., 2019; Jenkins et al., 2014; Stedman et al., 2017, fish: Crespel et al., 2011). Glucocorticoid concentrations also respond to artificial selection within a few generations (Evans et al., 2006; Pottinger & Carrick, 1999; Satterlee & Johnson, 1988; Touma et al., 2008), suggesting that this endocrine system can evolve under directional selection.

Alterations in environmental temperatures are one of the most prominent manifestations of the ongoing global climate change, both in yearly statistics and in the frequency of extreme events (IPCC, 2014). It is therefore critical to determine individual variation in glucocorticoid responses in wild populations to this key ecological factor, to appraise the impact of global warming on vertebrate populations and communities (Mentesana & Hau, in revision; Ruuskanen et al., 2021). To our knowledge, it is presently unknown whether individuals from free-living vertebrate populations differ in their glucocorticoid plasticity when experiencing natural fluctuations in ambient temperature. Addressing this question requires a reaction norm approach, which in its simplest form describes a linear relationship between an individual’s repeatedly measured phenotype (e.g., glucocorticoid concentrations) and an environmental or internal gradient (e.g., ambient temperature, Via et al., 1995). Using linear mixed models, phenotypic plasticity can be decomposed in two reaction norm parameters: the elevation (the average value of a trait in the average environment) and the slope of the response to the environment (the change in the trait across the gradient: its plasticity; Dingemanse et al., 2010; Nussey et al., 2007). Furthermore, these linear mixed models allow to test whether elevation and slope covary, i.e., whether links exist between these two reaction norm components that may constrain their variation (Dingemanse et al., 2010; Nussey et al., 2007).

Reaction norm analyses can be used to analyze glucocorticoid variation separately at two levels: baseline versus stress-induced concentrations. This distinction is important because at these two levels glucocorticoids bind to different genomic receptors and can exert divergent phenotypic effects (Koolhaas et al., 2011; Landys et al., 2006; Romero & Wingfield, 2016; Sapolsky et al., 2000). At low baseline levels, circulating glucocorticoid concentrations are thought to vary with predictable events like changes in weather or life history stage and to bind predominantly to the high-affinity mineralocorticoid receptor (Landys et al., 2006). In doing so, they promote plastic adjustments in behavioral and physiological traits prior to, during, and after these predictable challenges, for example changes in foraging behavior, offspring provisioning and metabolic rate (Landys et al., 2006). Whenever conditions change unpredictably and include major energetic or psychological challenges, glucocorticoids increase within a few minutes to reach high stress-induced concentrations, then also binding to the low-affinity glucocorticoid receptor (Koolhaas et al., 2011; Romero et al., 2009; Sapolsky et al., 2000; there also exists a membrane receptor with rapid, non-genomic actions in most vertebrates, e.g., Borski, 2000). One of the functions of stress-induced glucocorticoid levels is to support a switch to an ‘emergency state’, which can include mobilization of energy stores, re-allocation of energetic resources to processes that facilitate coping with the immediate challenge, and inhibition of behaviors like parental care that are not essential for surviving this challenge (Sapolsky et al., 2000; Wingfield et al., 1998). Stress-induced glucocorticoid levels are also thought to help the individuals to recover from and prepare for subsequent challenges (Romero & Wingfield, 2016; Sapolsky et al., 2000).

Recent reports suggested that individual differences in glucocorticoid reaction norm elevations and slopes exist in wild vertebrate populations (summaries: Guindre-Parker, 2020; Malkoc, Mentesana, et al., 2021; Taff & Vitousek, 2016). However, some of these studies determined glucocorticoid metabolites from excrements (feces: Guindre-Parker et al., 2019; urine: Sonnweber et al., 2018), which can be additionally influenced by individual differences in dietary, metabolic, digestive and other processes (reviews in: Dantzer et al., 2014; Gormally & Romero, 2020; Goymann, 2012). Furthermore, because glucocorticoid metabolites accumulate in urine and feces for hours to days before excretion, these studies cannot distinguish between patterns occurring at baseline versus stress-induced levels, or between events occurring at acute versus more distant time scales (e.g., Romero & Beattie, 2021). Other studies measured glucocorticoid concentrations in the blood along internal gradients of increasing age or breeding stage, but focused on aspects of the ‘stress-response’, i.e. the increase from baseline to stress-induced concentrations (Lendvai et al., 2015, Grant et al., 2020). Two captive studies on birds documented individual variation in baseline glucocorticoid plasticity, while two captive fish studies found individual differences only in glucocorticoid elevation (in water-borne hormone samples; Fürtbauer et al., 2015; Houslay et al., 2019). Findings in captivity can be of limited relevance in natural conditions because individuals may be chronically stressed and/or lack natural cues, which can bias glucocorticoid concentrations (Beaulieu, 2016; Calisi & Bentley, 2009; Cyr & Romero, 2009; Fusani et al., 2005).

We therefore conducted a 5-year study on a population of free-living great tits (*Parus major*) where we applied a reaction norm approach to determine within- and among-individual variation in glucocorticoid plasticity at baseline and stress-induced levels along natural fluctuations in ambient temperature. We repeatedly sampled adults of both sexes for circulating corticosterone concentrations (the main avian glucocorticoid) during a standardized phase of reproduction, the peak nestling provisioning phase. Using a series of general linear-mixed effect models (GLMMs) to analyze baseline and stress-induced corticosterone variation separately, we tested the following predictions:

1. Population-level relationships with ambient temperature: baseline and stress-induced corticosterone concentrations are higher at lower ambient temperatures. This is a common pattern among endotherms (summaries: de Bruijn & Romero, 2018; Jessop et al., 2016; Ruuskanen et al., 2021; Wingfield & Ramenofsky, 2011), and likely results from more energy being expended on thermoregulation and increased foraging activity at lower temperatures (e.g., Rubalcaba & Jimeno, 2022). We also predicted that baseline and stress-induced corticosterone concentrations are more strongly associated with short-term (temperature at capture) than with longer-term temperature measurements (mean temperatures on capture day or on days preceding capture) because parental great tits should track short-term temperature changes as these affect their own energy requirements as well as their foraging effort.
2. Among-individual differences in *average* corticosterone concentrations: individuals differ in average baseline and stress-induced corticosterone concentrations. To test this prediction, we included individual ‘identity’ (ID) as a random factor in our models, expecting it to explain a significant amount of the variation in corticosterone concentrations. In recent meta-analyses across vertebrate taxa, the total phenotypic variation explained by among-individual differences (‘individual repeatability’) was between 18-23% for baseline and around 38% for stress-induced glucocorticoid concentrations (meta-analyses: Schoenemann & Bonier, 2018; Taff et al., 2018).
3. Within-individual *changes* in corticosterone concentrations with temperature: an individual bird has higher corticosterone levels when being sampled at lower than at higher ambient temperatures. In great tits, the thermoneutral zone (the range of environmental temperatures at which individuals do not expend energy on thermoregulation) ranges between 15-30°C (Broggi et al., 2005; Malkoc, Casagrande, et al., 2021). During the breeding season, birds from our population commonly experience temperatures below the thermoneutral zone (Fig. S1) and then should increase corticosterone concentrations (see prediction 1). Using the ‘within-individual centering’ approach (van de Pol & Wright, 2009), we tested the hypothesis that single individuals indeed plastically change corticosterone concentrations with ambient temperature and that population-level patterns were not caused by sampling different individuals at different temperatures. Studies documenting within-individual glucocorticoid plasticity to natural temperature variation are exceedingly rare (in captive birds: Jimeno et al., 2018b).
4. Among-individual differences in corticosterone *plasticity* in response to temperature: individual great tits differ in their corticosterone reaction norm slope to the temperature gradient, with some individuals showing a greater plasticity (steeper slope) than others. To test this, we added a random factor for the interaction between ID and temperature to the model built to test prediction 2. A recent captive study on male house sparrows (*Passer domesticus*) has indicated that individuals differ in the baseline corticosterone slope to experimental heat exposure (Baldan et al., 2021).
5. Among-individual covariation between *elevation* and *slope*: we explored whether these two reaction norm attributes covary for baseline or stress-induced corticosterone concentrations. A trend for a covariation was observed in captive Trinidadian guppies (*Poecilia reticulata*) (Houslay et al., 2019), but thus far has rarely been tested (summarized by Guindre-Parker, 2020).
6. Population-level, among- and within-individual covariation between baseline and stress-induced corticosterone concentrations: baseline and stress-induced corticosterone concentrations covary positively at the population and within-individual levels. This prediction is based on earlier findings that a great tit individual sampled at a particular time with higher baseline corticosterone also had higher stress-induced concentrations (within-individual covariation, Baugh et al., 2014), but great tits with higher average baseline corticosterone did not have a higher average stress-induced concentrations (no among-individual covariation, Baugh et al., 2014; similar findings: Jenkins et al., 2014; Stedman et al., 2017; evidence for genetic covaration: Béziers et al., 2019).

## Results

### Population-level corticosterone responses to temperature, and individual differences in average concentrations

Variation in baseline and stress-induced corticosterone was best explained by ambient temperature at the moment of capture (based on DIC model comparisons, Supplementary Material Table S1-2). Baseline and stress-induced corticosterone were both negatively related to temperature at capture (Table 1, Figure 1), i.e., baseline and stress-induced corticosterone levels were higher when temperatures at capture were lower. This significant effect of temperature at capture was somewhat stronger for stress-induced corticosterone than for baseline corticosterone (Table 1, Figure 1), indicating that stress-induced corticosterone might be even more conditioned by local temperatures than baseline concentrations. Furthermore, as expected from ample previous work including on great tits (Baugh et al., 2013; Romero & Reed, 2005; Small et al., 2017; Wingfield et al., 1982), baseline, but not stress-induced corticosterone, was influenced by the time needed to take a blood sample, with longer times needed to obtain a sample being associated with higher concentrations of baseline corticosterone. We also found large year differences for both hormonal traits, but no sex differences (Table 1). Both hormonal traits were repeatable: individual differences in average trait expression explained 17% of the total population variance in baseline corticosterone, and 26% in stress-induced corticosterone (Table 1).

**Table 1.** Sources of phenotypic variation in baseline and stress-induced corticosterone concentrations. Both variables were divided by 1000 and modelled following a Gaussian error distribution. Estimates of fixed (β) and random (σ2) parameters are shown as posterior modes with 95% Credible Intervals (CI).

| Fixed effects | Baseline corticosterone |  | Stress-induced corticosterone |  |
| --- | --- | --- | --- | --- |
| | $\beta$ | 95% CI | $\beta$ | 95% CI |
| Intercept | 6.817 | [5.935 , 7.696] | 23.904 | [21.071 , 26.323] |
| Sex [female] | 0.696 | [-0.064 , 1.489] | 2.308 | [-0.038 , 4.726] |
| Year [2016] | -1.602 | [-2.663 , -0.622] | -6.404 | [-9.673 , -3] |
| Year [2017] | -1.891 | [-2.91 , -0.826] | -1.634 | [-4.887 , 1.571] |
| Year [2018] | -1.252 | [-2.329 , -0.095] | 7.52 | [3.882 , 11.196] |
| Year [2019] | -3.452 | [-4.641 , -2.243] | -0.604 | [-4.06 , 2.941] |
| Bleeding time (scaled) | 1.022 | [0.609 , 1.404] | 0.561 | [-0.644 , 1.799] |
| Temperature at capture (scaled) | -0.678 | [-1.006 , -0.275] | -2.78 | [-3.939 , -1.716] |
| Random effects | $\sigma^2$ | | $\sigma^2$ | |
|  |  | 95% CI |  | 95% CI |
| Individual | 2.141 | [0.05 , 4.119] | 32.8 | [13.372 , 53.256] |
| Residual | 10.42 | [8.047 , 12.824] | 93.496 | [74.123 , 115.069] |
| Repeatability | 0.169 | [0.01 , 0.313] | 0.258 | [0.122 , 0.405] |

**Figure 1.**
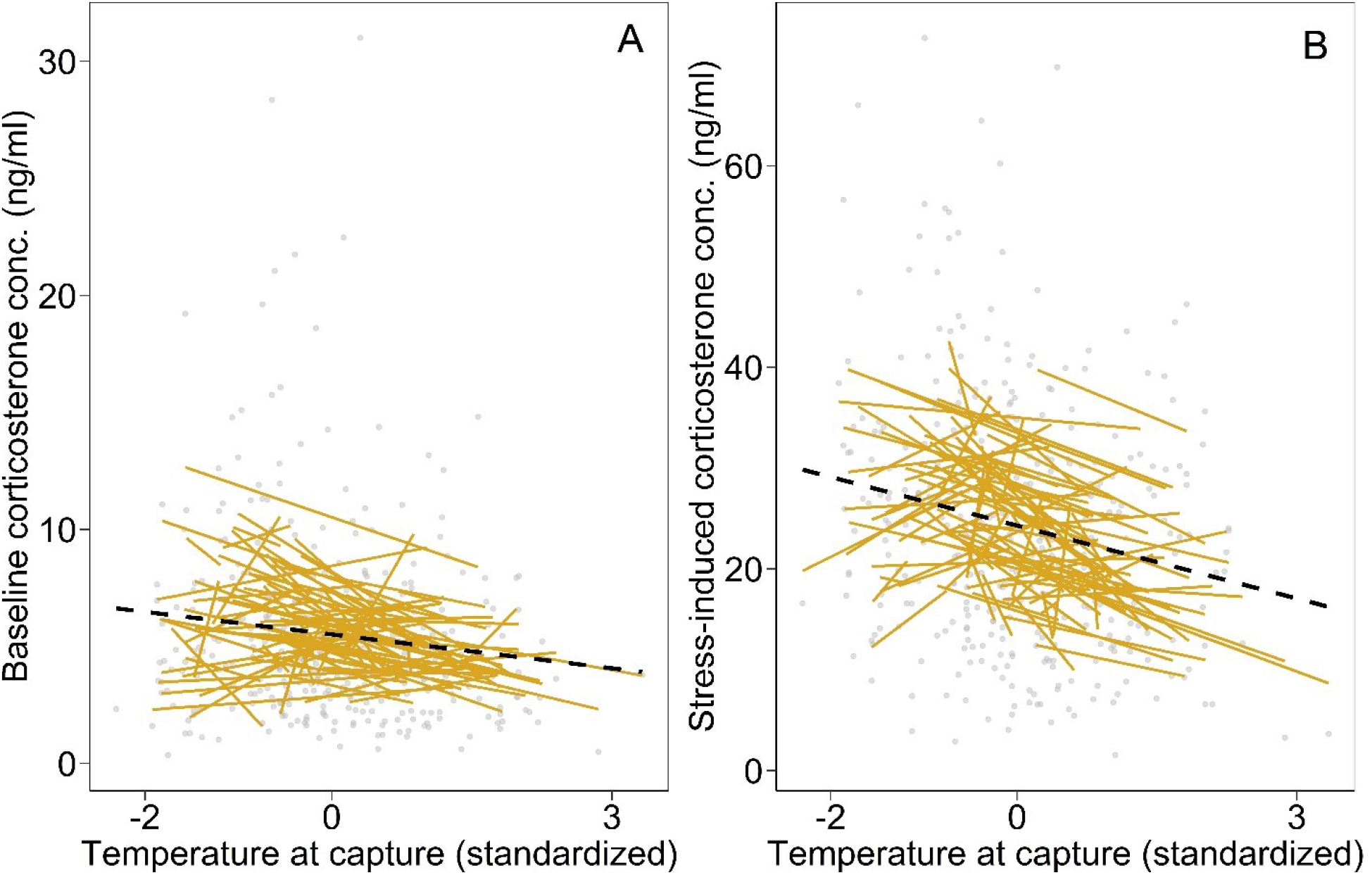
Reaction norm model predictions from two random regression models of baseline (A) and stress-induced (B) corticosterone in response to temperature at capture (mean centered and variance standardized), respectively. Each yellow line represents a single individual, the dashed black line represents the population-level response to temperature, and grey dots represent the raw phenotypic data

### Within-individual changes in response to temperature

Both baseline and stress-induced corticosterone showed significant within-individual variation in response to ambient temperature (βwithin-ID temperature effect; Table 2). This indicates that single individual great tits plastically adjusted their hormonal levels from one sampling event to another in response to variation in temperature (within-individual plasticity). There was also between-subject variation (βbetween-ID temperature effect) in baseline and stress-induced corticosterone (although the effect for baseline corticosterone is slightly overlapping zero, Table 2). This implies that those individuals, that on average experienced lower temperatures across the 5-year sampling period, tended to have on average higher baseline and stress-induced corticosterone levels. There was no significant difference between the within- and between-subject effects for either baseline (posterior mode β_B_–β_W_ = 0.567, 95%CI = -0.114, 1.289) or stress-induced corticosterone (posterior mode β_B_–β_W_ = -0.673, 95%CI = -3.073 1.231), suggesting that a single mechanism explains the observed patterns: plastic responses of individuals to temperature variation underlie the population-level relationship with temperature for both hormonal traits.

**Table 2.**
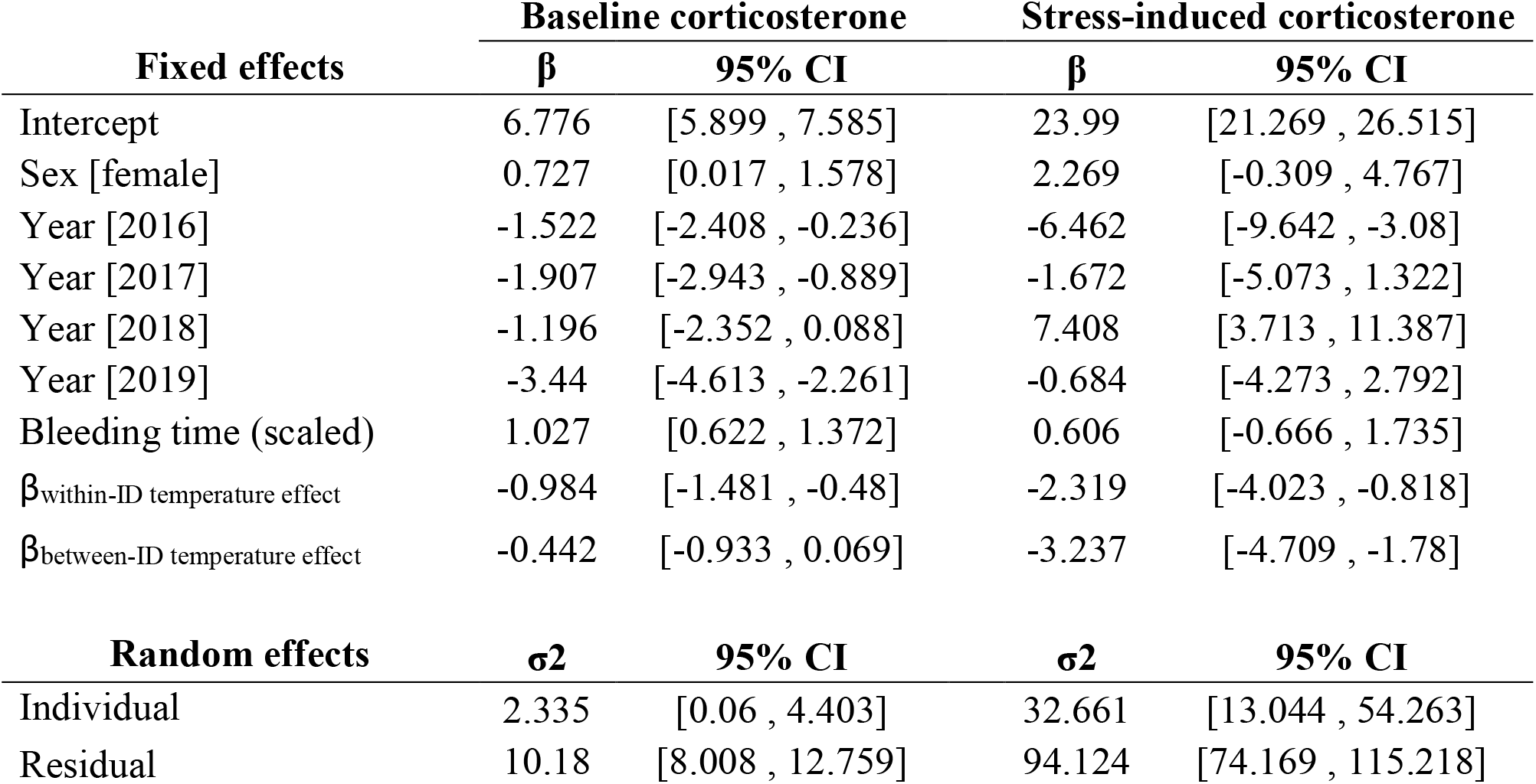
Estimates of between- and within-individual effects of temperature at capture on baseline and stress-induced corticosterone modelled using a “within-subject centering” approach. Estimates of fixed (β) and random (σ2) parameters are shown as posterior modes with 95% Credible Intervals (CI).

| Fixed effects | Baseline corticosterone |  | Stress-induced corticosterone |  |
| --- | --- | --- | --- | --- |
| | $\beta$ | 95% CI | $\beta$ | 95% CI |
| Intercept | 6.776 | [5.899 , 7.585] | 23.99 | [21.269 , 26.515] |
| Sex [female] | 0.727 | [0.017 , 1.578] | 2.269 | [-0.309 , 4.767] |
| Year [2016] | -1.522 | [-2.408 , -0.236] | -6.462 | [-9.642 , -3.08] |
| Year [2017] | -1.907 | [-2.943 , -0.889] | -1.672 | [-5.073 , 1.322] |
| Year [2018] | -1.196 | [-2.352 , 0.088] | 7.408 | [3.713 , 11.387] |
| Year [2019] | -3.44 | [-4.613 , -2.261] | -0.684 | [-4.273 , 2.792] |
| Bleeding time (scaled) | 1.027 | [0.622 , 1.372] | 0.606 | [-0.666 , 1.735] |
| $\beta_{\text{within-ID temperature effect}}$ | -0.984 | [-1.481 , -0.48] | -2.319 | [-4.023 , -0.818] |
| $\beta_{\text{between-ID temperature effect}}$ | -0.442 | [-0.933 , 0.069] | -3.237 | [-4.709 , -1.78] |
| Random effects | $\sigma^2$ | | $\sigma^2$ | |
| | $\sigma^2$ | 95% CI | $\sigma^2$ | 95% CI |
| Individual | 2.335 | [0.06 , 4.403] | 32.661 | [13.044 , 54.263] |
| Residual | 10.18 | [8.008 , 12.759] | 94.124 | [74.169 , 115.218] |

### Among-individual differences in corticosterone plasticity and elevation-slope covariance

We tested whether individuals differed in their plastic responses to temperature (i.e., for among-individual variation in reaction norms). For both baseline and stress-induced corticosterone, DIC model comparison showed that the best-fitting model included a random factor for the interaction between individual identity and temperature at capture. This model also allowed residual variation to differ among the 5 years of sampling (i.e., was a model with heterogeneous residual variance; Table 3). The results from this model imply that individuals differ in average baseline and stress-induced corticosterone at the population’s average temperature (i.e., among-individual variation in reaction norm elevation) and that individuals also differed in baseline and stress-induced corticosterone plasticity, with some individuals responding more strongly to temperature changes than others (i.e., among-individual variation in reaction norm slopes; Table 4, Figure 1). For baseline corticosterone, we found a negative covariation of elevation and slope, indicating that individuals with lower *or* higher average concentrations of baseline corticosterone had steeper responses to the temperature gradient (“fanning-in” pattern, Figure 1).

**Table 3.** DIC model comparison of a series of univariate mixed-effects models for baseline and stress-induced corticosterone including and excluding an individual random slope term for temperature at capture, and/or heterogeneous residuals. Heterogeneous residual structure was modelled in five blocks (one per year of sampling).

| Trait | Random effects | Residuals | DIC | ΔDIC |
| --- | --- | --- | --- | --- |
| Baseline corticosterone | ID: random intercepts | homogeneous | 2077.69 | 309.04 |
|  | ID: random intercepts | heterogeneous (5) | 1904.543 | 135.89 |
|  | ID: random intercepts & slopes | homogeneous | 2069.675 | 301.02 |
|  | ID: random intercepts & slopes | heterogeneous (5) | 1768.655 | 0.00 |
| Stress-induced corticosterone | ID: random intercepts | homogeneous | 2859.398 | 47.21 |
|  | ID: random intercepts | heterogeneous (5) | 2824.747 | 12.56 |
|  | ID: random intercepts & slopes | homogeneous | 2859.385 | 47.20 |
|  | ID: random intercepts & slopes | heterogeneous (5) | 2812.189 | 0.00 |

**Table 4.** Reaction norm results from a random regression model of baseline and stress-induced corticosterone as a function of temperature at capture. Estimates of fixed (β) and random (σ2) parameters are shown as posterior modes with 95% Credible Intervals (CI).

| Fixed effects | Baseline corticosterone |  | Stress-induced corticosterone |  |
| --- | --- | --- | --- | --- |
| | $\beta$ | 95% CI | $\beta$ | 95% CI |
| Intercept | 6.83 | [5.593 , 8.16] | 24.03 | [21.224 , 26.999] |
| Sex [female] | 0.69 | [0.025 , 1.226] | 2.43 | [0.194 , 4.912] |
| Year [2016] | -1.67 | [-3.055 , -0.364] | -6.43 | [-10.053 , -2.803] |
| Year [2017] | -1.91 | [-3.263 , -0.548] | -1.33 | [-4.588 , 2.077] |
| Year [2018] | -1.24 | [-2.632 , 0.302] | 7.35 | [3.26 , 11.122] |
| Year [2019] | -3.38 | [-4.78 , -2.097] | -0.74 | [-3.805 , 2.544] |
| Bleeding time (scaled) | 1.00 | [0.672 , 1.296] | 0.75 | [-0.411 , 1.902] |
| Temperature at capture (scaled) | -0.62 | [-0.856 , -0.353] | -2.39 | [-3.545 , -1.284] |
| Random effects | $\sigma^2$ | lower 95%CI | $\sigma^2$ | lower 95%CI |
| **Individual** |  |  |  |  |
| Intercept | 1.90 | [0.804 , 3.061] | 41.87 | [22.058 , 60.442] |
| Linear Slope | 0.29 | [0.001 , 0.658] | 3.60 | [0.009 , 10.882] |
| Intercept–slope covariance | -0.47 | [-0.957 , 0.012] | -2.88 | [-9.884 , 2.937] |
| Intercept–slope correlation | -0.65 | [-0.987 , -0.207] | -0.28 | [-0.895 , 0.352] |
| Residual 1 | 30.88 | [20.679 , 41.038] | 111.31 | [66.323 , 159.672] |
| Residual 2 | 6.21 | [3.768 , 8.923] | 102.74 | [65.293 , 141.702] |
| Residual 3 | 5.96 | [3.96 , 8.473] | 81.00 | [45.969 , 112.406] |
| Residual 4 | 6.42 | [3.671 , 9.286] | 102.66 | [58.087 , 153.517] |
| Residual 5 | 1.15 | [0.277 , 2.318] | 15.26 | [0.281 , 38.185] |

### Covariation between baseline and stress-induced corticosterone concentrations

We found a significant positive correlation between baseline and stress-induced corticosterone concentrations at the population level (Table 5), indicating that higher values of baseline corticosterone were associated with higher values of stress-induced corticosterone. When we statistically tested whether this phenotypic correlation is also present among and within individuals, we observed a positive correlation at both levels (Table 5). The positive correlation among individuals indicates that birds with on average higher baseline corticosterone concentrations also had on average higher stress-induced corticosterone than others. The positive within-individual correlation means that at the same sampling event, when a bird displayed high baseline corticosterone concentrations, it also displayed high stress-induced corticosterone concentrations.

**Table 5.** Results from a bivariate mixed-effects model of the correlations between baseline and stress-induced corticosterone at the phenotypic, among- and within-individual levels. Estimates represent the posterior modes for the correlation (*r*) and associated 95% Credible Intervals (CI).

| Phenotypic |  | Among-individual |  | Within-individual |  |
| --- | --- | --- | --- | --- | --- |
| $r$ | 95% CI | $r$ | 95% CI | $r$ | 95% CI |
| 0.326 | [0.223 , 0.407] | 0.35 | [0.144 , 0.738] | 0.25 | [0.102 , 0.388] |

## Discussion

In this study, we applied a reaction norm approach to data from free-living individuals repeatedly sampled for circulating glucocorticoid concentrations across five years. We obtained support for our prediction that individuals plastically change circulating baseline and stress-induced corticosterone concentrations in response to the ambient temperature they experienced in each year during the peak nestling provisioning phase (Figure 1, Table 2). Additionally, we found that within-individual plasticity explained the negative population-level relationship observed between both corticosterone traits and ambient temperature. Our reaction norm approach revealed that individuals not only differ in average corticosterone (i.e., in the reaction norm elevation of baseline and stress-induced concentrations), but also in the slope of their reaction norm: some individuals plastically adjusted baseline and stress-induced corticosterone concentrations more strongly to ambient temperature than others (Figure 1, Tables 3 & 4). These findings provide unique evidence that plastic endocrine responses to a key ecological factor vary among and within individual free-living great tits. Thus, if individual variation in glucocorticoid plasticity harbored heritable variation (e.g., Bairos-Novak et al., 2018; Jenkins et al., 2014) and was under selection, it would provide the basis for adaptive evolution to occur in a conserved higher-order mechanism that enables vertebrates to adjust their phenotype to fluctuations in climatic conditions. Altogether, our findings are particularly relevant in light of the ongoing global changes in climate, which are expected to increase the frequency of temperature extremes world-wide (IPCC, 2014).

Our prediction that corticosterone concentrations are higher at lower ambient temperatures was confirmed, but the mechanism underpinning this pattern is not yet clear. In our study, temperatures at capture ranged between ∼4-27°C (Figure S1). Assuming that the thermoneutral zone of our great tit population is between 15°C and 30°C (Broggi et al., 2005; Malkoc, Casagrande, et al., 2021), we collected two thirds of the baseline and stress-induced corticosterone samples at temperatures below the great tits’ thermoneutrality. The thermoneutral zone is typically determined in a laboratory situation and only insufficiently reflects thermoregulatory challenges and processes that animals face under natural conditions (e.g., Mitchell et al., 2018). However, even though the exact temperatures and pathways that affect glucocorticoid concentrations are still unclear, two scenarios are conceivable. First, cooler ambient temperatures may have induced increases in corticosterone concentrations in great tit individuals directly via sensory information collected by thermoreceptors and integrated in hypothalamic brain areas (reviewed in: McKechnie, 2022; Ruuskanen et al., 2021; Yahav, 2015). Second, an indirect pathway could involve metabolic rate, which likely was higher in great tit individuals sampled at cooler than at warmer temperatures (e.g., Broggi et al., 2004), because of their need to generate heat through shivering and non-shivering thermogenesis (McKechnie, 2022). Additionally, individuals caught at lower temperatures likely also showed a greater foraging activity to provision their 8-10 day old young with insect food that is harder to find and less abundant at cold temperatures (e.g., Avery & Krebs, 1984; Scholl et al., 2016) – which can also increase metabolic rate (Nilsson, 2002; Wiersma & Tinbergen, 2003). Metabolic rate in turn can be positively associated with glucocorticoid concentrations within individuals (Jimeno et al., 2017, 2018a). At present we cannot exclude that other factors associated with ambient temperature, for example precipitation, wind or barometric pressure contributed to within-individual corticosterone plasticity (de Bruijn & Romero, 2018; Wingfield & Ramenofsky, 2011).

Great tit individuals showed a negative relationship in both baseline and stress-induced corticosterone with ambient temperature (Table 4), suggesting that this ecological variable affects the functioning of this endocrine system when dealing with predictable homeostatic processes as well as its response to additional unpredictable challenges, as simulated by the capture-restraint protocol. We considered it important to analyze baseline and stress-induced glucocorticoid concentrations separately, as they are known to be stimulated by different types of environmental information (predictable versus unpredictable challenges), mostly act via different genomic receptors (mineralocorticoid versus glucocorticoid), and support divergent phenotypic effects (e.g., Koolhaas et al., 2011; Landys et al., 2006; Romero & Wingfield, 2016; Sapolsky et al., 2000). The positive covariation between baseline and stress-induced corticosterone at the population, among-individual, and within individual level (Table 5) does not contrast with this view. A likely scenario is that ambient temperature affects an individual’s allostatic load or reactive homeostasis (i.e., the cumulative challenges, energetic or otherwise, that an individual has to cope with at one point in time; McEwen & Wingfield, 2003; Romero et al., 2009), and this change in state could influence both traits. At present there is no evidence for genetic covariation between the two glucocorticoid traits in free-living songbirds (Jenkins et al., 2014; Stedman et al., 2017, but for non-songbirds see: Béziers et al., 2019).

As expected, individuals in our study also differed in average baseline and stress-induced concentrations (Table 1, Figure 1). Our corticosterone repeatability estimates (baseline: 17%; stress-induced: 26%) are within the range of those reported in recent meta-analyses (across vertebrate taxa, baseline: 18-23%; stress-induced: 37-38%, (Schoenemann & Bonier, 2018; Taff et al., 2018). These individual differences in average glucocorticoid concentrations align well with heritability estimates of those traits in studies on free-living bird populations, where additive genetic effects explained less than 20% of the variance in baseline and more than 30% in stress-induced concentrations, respectively (barn swallows: Jenkins et al., 2014; tree swallows Stedman et al., 2017, barn owls: Béziers et al., 2019).

The presence of individual-by-environment interactions (I x E) in our great tit population, with some individuals showing weak and others pronounced plasticity in baseline and stress-induced corticosterone concentrations along a temperature gradient (Figure 1, Table 4), is intriguing from a proximate viewpoint. In principle, individual differences in plasticity could arise from two processes: a) long-term individual differences that are based on genetic information, maternal or other early developmental effects, b) more short-term variation in an individual’s state or c) both (e.g., Dingemanse & Wolf, 2013; Sih et al., 2015). Both long- and short-term processes could generate among-individual differences in plasticity via variation in, for example, the perception and neuro-endocrine processing of environmental temperature, the capacity to synthesize glucocorticoids (including the sensitivity of the hypothalamo-pituitary-axis to releasing hormones), the binding of glucocorticoids to carrier molecules in the blood, their enzymatic deactivation, and differences in receptor densities (Hau et al., 2016; Taff & Vitousek, 2016; Wingfield, 2013). In future studies, the contribution of genetic components to among-individual differences in corticosterone plasticity could be quantified in statistical models that include information on pedigrees, while simultaneously quantifying maternal and early developmental contributions using correlative data or experimental approaches like cross-fostering (Taff & Vitousek, 2016; Wada & Sewall, 2014; Wilson et al., 2010). Variation in an individual’s short-term state can also underpin individual differences in corticosterone plasticity: although we sampled great tits during a standardized parental phase, individuals could differ in various aspects, including body condition, health status, territory or partner quality, reproductive investment and age (the sexes did not differ in this study). Many of these variables determine an individual’s energy intake and/or usage and circulating glucocorticoid concentrations (McEwen & Wingfield, 2003; Romero et al., 2009). Additionally, if glucocorticoid plasticity was energetically costly, perhaps via its effects on phenotypic traits like locomotor and foraging activity, an individual’s energetic state may determine its degree of plasticity (Malkoc, Mentesana, et al., 2021). However, in a recent study on captive house sparrows (*Passer domesticus*), statistically accounting for body mass plasticity did not explain the existence of individual corticosterone plasticity (Baldan et al., 2021; Lendvai et al., 2014). In our study birds, we did not determine individual differences in energy intake or usage like metabolic rate.

However, captive zebra finches show individual differences in the plasticity of metabolic rate to changes in ambient temperature (Briga & Verhulst, 2017). We therefore consider this to be a likely pathway to generate individual differences in glucocorticoid plasticity in our great tit population. Technologies like the measurement of mitochondrial metabolic rate in red blood cells can help test this hypothesis in free-living great tits (Malkoc, Casagrande, et al., 2021; Stier et al., 2017).

In individuals from our study population, the elevation and slope of reaction norms in baseline corticosterone were correlated, suggesting that these two features of the endocrine response to environmental temperature in free-living great tits are linked (Table 4). The fanning-in pattern evident in baseline corticosterone slopes towards higher ambient temperatures (Figure 1a) suggests that individuals with both higher and lower elevation had steeper slopes, while individuals with medium elevation had more shallow slopes. At present, we cannot determine the reason for this complex association between elevation and slope in baseline corticosterone of wild great tits. We can, however, speculate that differences in the individuals’ energetic or health state may play a role. Glucocorticoids can have context-, state- and tissue-dependent effects: they can mobilize fat reserves in adipose tissue but also increase fat storage in the liver (reviewed in: Landys et al., 2006). In migratory birds, concentrations of corticosterone, mRNA expression of glucocorticoid receptors and re- and deactivating enzymes can differ between seasonal states and muscle types (e.g., Pradhan et al., 2019). Thus, if individuals with a low baseline corticosterone elevation were in an anabolic state, they could adjust corticosterone concentrations to ambient temperature fluctuations differently than individuals in a catabolic state (for example those with medium or high corticosterone elevation). This scenario could be tested by carefully assessing an individual’s energetic state (e.g., via morphometric, or blood and tissue composition measures), ideally coupled with experimental manipulation of energetic challenges, for example by increasing the workload of parents by clipping wing feathers (Casagrande & Hau, 2018) or manipulating brood size (Bonier et al., 2011). From an evolutionary viewpoint, the covariation between elevation and slope is important because it suggests that these two attributes of baseline corticosterone cannot be studied in isolation (e.g., Malkoc, Mentesana, et al., 2021). It will be rewarding in future studies to investigate the causes underlying this pattern, as well as its functional consequences.

The current study is one of only few to analyze individual differences in endocrine reaction norms in a free-living vertebrate population (Grant et al., 2020; Guindre-Parker et al., 2019; Hsu et al., 2019; Lendvai et al., 2015; Mentesana et al., 2019; Sonnweber et al., 2018). Repeatedly capturing and collecting blood samples from free-living individuals during a standardized time of year is logistically challenging (Guindre-Parker, 2020; Malkoc, Mentesana, et al., 2021). Of the repeatedly sampled individuals in our 5-year study, we were able to collect two samples from more than 60 individuals (68 for baseline, and 62 for stress-induced corticosterone) and over 30 individuals (32 for baseline, and 31 for stress-induced corticosterone) could be sampled 3 or more times. Sampling an individual at least twice is the minimum requirement for reaction norm studies, but by retaining individuals that were sampled only once (140 for baseline and 144 for stress-induced) in our statistical models, we improved the estimation of the phenotypic variance at the population level (Martin et al., 2011; van de Pol, 2012).

## Conclusions

Large fluctuations in environmental temperature during the breeding season are common for our great tit population (Figure S1), perhaps explaining the existence of corticosterone plasticity to this important ecological gradient as well as the among-individual variation in slopes. It will now be important to connect individual variation in glucocorticoid reaction norm components to the expressed phenotype of an individual, for example reproductive investment like provisioning rates, reproductive decisions like nest desertion and fitness-relevant traits like reproductive success, recruitment and survival (Bonier & Martin, 2016; Cox et al., 2016; Dantzer & Swanson, 2017; Hau et al., 2016; Malkoc, Mentesana, et al., 2021; Taff & Vitousek, 2016; Wada & Sewall, 2014). Future work should also aim at elucidating the pathways through which environmental temperatures induce glucocorticoid plasticity in birds and other vertebrates. Finally, this line of work will be important for evaluating the consequences of climate change for wild vertebrate populations. Although climate change is predicted to increase global temperatures, it is also expected to increase the frequency of extreme temperature events (IPCC, 2014). Bird populations that have advanced their breeding phenology in response to a warming climate now experience cold snaps more frequently during the breeding season (Shipley et al., 2020). The work presented here has documented the existence of individual differences in corticosterone responses to acute changes in ambient temperature. Directional selection on such individual differences can lead to divergent corticosterone responses to cold exposure in poultry within a few generations (Brown & Nestor, 1973). Collectively, these data provide an excellent basis for investigating how wild vertebrate populations may cope with future changes in ambient temperature regimes.

## Methods

### Field work

Adult great tits were studied in a nest box population in the Ettenhofer Holz, Upper Bavaria, Germany from early May to early July of 2015 through 2019. Adult great tits were captured when entering the nest box to feed their nestlings using a remote-controlled or mechanical trap, or by blocking the entrance hole with a bird bag when nestlings were about 8 days old (hatching = day 0). Captures were restricted to the morning hours between 7:00h and 14:00h to minimize effects of diel variation in corticosterone concentrations. We aimed at catching both parents on the same day, but if unsuccessful tried to capture the missing parent within the next two days. Also, we avoided capturing parents during inclement weather conditions by either advancing capture by one day or delaying it by maximally three days. Therefore, adult birds were caught when chicks were between 7 and 11 days old, which is when provisioning rates are very high and/or peak (Sanz & Tinbergen, 1999). Most birds were captured only once every year and only few individuals (29 out of 240) were captured twice within a year. Nests were checked during egg laying and incubation every 1-2 days to determine laying and hatching dates.

### Temperature recordings

Temperature at capture was recorded using a handheld thermometer (BASETech Thermometer E0217, accuracy ±1°C). Whenever temperature could not be directly recorded, we used data from a weather station (HOBO weather station with S-THB-M002 12 Bit Temperature and relative humidity sensor, logging ambient temperature every 30min) located within our study area. In that case, we used the temperature of the 30min increment closest to the time of capture. Mean, maximum or minimum temperatures used in the statistical analyses were also calculated from the weather station data.

### Blood sampling and hormone analyses

Blood was sampled for baseline corticosterone within about 3 min of blocking of the entrance hole of the nest by pricking the wing vein with a 26 g needle and collecting the blood in heparinized microcapillaries. The overall (mean ± SD) time to finish blood sampling for baseline corticosterone was 2.1 ± 0.58 min, with a range of 0.92 to 4 min. After the bleeding had stopped and a 1 min video recording of the bird was taken, it was put in an opaque cotton bag until banding. Biometric measurements (mass, tarsus and wing length, muscle and fat scores) and a second blood sample for stress-induced corticosterone were taken about 30 min after the initial capture (mean ± SD: 30.1 ± 2.88 min, range: 19.75 to 45 min). Blood samples were kept on ice until centrifugation in our laboratory at 4°C. The plasma fraction was removed with a pipette and stored at -20°C until analysis. Following the second blood sample, we returned the bird into the nest box.

Samples were analyzed for corticosterone concentrations using commercially available enzyme-linked immunosorbent assays, following a double liquid-liquid extraction with diethyl ether (Baugh et al., 2012; Ouyang et al., 2011). The reconstituted samples were assayed in duplicate. Samples from 2015 were analyzed with the ENZO Life Sciences Corticosterone ELISA kit (ADI-900-097) (for a validation for great tit plasma samples see Ouyang et al., 2011). From 2016 onwards, all samples were analyzed with Arbor Assays DetectX® Corticosterone Enzyme Immunoassay Kit (K014-H5). Given that we analyzed hormone samples in the first year (2015) with immunoassays from a different company than in the following years (2016-2019), we included ‘year’ as a fixed factor in all our models (see below). We could not model differences across ELISA manufacturers independently from variation among years, because differences in the year 2015 may result from both assay manufacturer and overall environmental conditions when samples were collected.

The Arbor Assays kit was validated for its use with great tit plasma by assessing parallelism and accuracy through the recovery of spiked plasma samples. We assessed parallelism by serially diluting three great tit plasma pool samples (adult baseline, adult stress-induced, nestlings stress-induced) which were parallel to the corticosterone standard curve in all three pool samples. Mean recovery for the three spiked pool samples was 108%, indicating no matrix interference with analyte detectability.

Control samples were also included in each assay batch and taken through the entire extraction and analysis procedure like great tit plasma samples. The positive controls consisted of stripped chicken plasma to which we added a known amount of a 100 000 pg/ml corticosterone stock solution for the final concentration. Negative controls containing only ultrapure water were included as the first and last sample of each extraction batch to detect possible contamination. For the double liquid-liquid extraction of samples and positive controls, 1-15µl of plasma were added to 200µl of HPLC-grade water in glass reagent tubes. No plasma was added to the ultrapure water for negative controls. Then, 1ml diethyl ether (Merck Product No. 1.00921) was added and samples were vigorously shaken for 10min. The phases were left to separate for 10 min, then each tube was placed in a dry ice bath with 99% ethanol to freeze the water phase, and the diethyl ether supernatant was decanted into a new glass tube. Each sample was thawed and frozen again, as some samples did not fully separate before. The additional ether phase was added to the respective sample glass tube. After the water phase had thawed, 1ml diethyl ether was added to each glass tube and the whole process was repeated a second time. The second supernatant was combined with the first supernatant. Then, the diethyl ether was evaporated under a constant nitrogen air stream in a dry heating block set to 37°C for ∼5min until completely dry. Samples were reconstituted with the assay buffer provided with the ELISA assay kits, briefly vortexed, covered with parafilm, and left to equilibrate at 4°C overnight until assayed the next day. The average extraction recovery for this procedure in our laboratory is around 90% (Baugh et al., 2012; Ouyang et al., 2011).

The reconstituted samples were assayed in duplicate, and all samples from the same individual in one year (baseline and stress induced, repeated samplings) were run on one plate. Baseline samples were assayed at a dilution of 1:10-1:20, and stress-induced samples at a dilution of 1:40-1:80, depending on the plasma volume available. Final corticosterone concentrations of samples were multiplied with the respective dilution factor. To assess assay precision, intra- and inter-assay coefficients of variation were calculated using the positive control samples. Because our positive control samples undergo the extraction procedure just like the actual samples, our inter-assay coefficient of variation incorporates the variation due to extraction efficiency and performance of each assay plate.

For ENZO assay plates, one control sample was measured in duplicate twice on each plate and used to calculate intra- and inter-assay variation. The absolute corticosterone concentration for this control sample was 25360 pg/ml, and it was measured at a 1:40 dilution, which corresponded to around 52% binding and an average concentration of 634 pg/ml. Inter-assay variation was 16% across eight plates. The mean intra-assay coefficient of variation calculated from the two measurements of the control sample on each plate was 13%.

For Arbor Assay plates, two control samples were measured in duplicate twice on each plate and were used to calculate intra- and inter-assay variation. One control sample with an absolute corticosterone concentration of 5550 pg/ml was measured at a 1:30 dilution factor, which corresponded to around 77% binding and an average concentration of 185pg/ml. Baseline samples at 1:10-1:15 dilutions were measured in this range of the assay. The second control sample with an absolute corticosterone concentration of 14700 pg/ml was measured at a 1:19 dilution factor, which corresponded to around 45% binding, with an average concentration of 782 pg/ml. Stress-induced samples were measured in this range of the assay. Inter-assay variation across 37 plates was 8% for the control measured at 45% binding, and 17% for the control measured at 77% binding. The mean intra-assay variation coefficient calculated for the two measurements of the same control sample on each plate was 9% for the control measured at 77% binding, and 4% for the control measured at 45% binding.

The detection limit of each plate was calculated as the mean optical density of the maximum binding (Zero, B_0_) duplicate wells minus two times the standard deviation between the duplicates. The corresponding concentration was determined against the plate’s standard curve. For samples in which the concentration was below the respective plate’s detection limit, the concentration given by the reader software was substituted with the concentration of the detection limit.

### Statistical analyses

For baseline corticosterone, we compiled a dataset of 389 measurements from 240 individuals. Among them, 140 individuals had one measurement, 68 individuals had two measurements in different years, 19 had three, 9 had four, and 4 individuals had five. For stress-induced corticosterone, the dataset contained 376 measurements from 237 individuals. Among them, 144 individuals had one measurement, 62 individuals had two measurements in different years, 19 had three, 9 had four, and 3 individuals had five.

#### Population-level corticosterone association with temperature measures

We ran a series of univariate general linear mixed-effects models to identify the critical temperature associated with baseline and stress-induced corticosterone in this population of great tits. We tested different temperature measures: temperature at the moment of capture, and mean, minimum and maximum temperatures on the day of capture, on the day prior to capture and on the three days prior to capture. All models included the fixed and random effects described below. We then used Deviance Information Criterion (DIC) model comparison to identify the best model (the one with the lowest DIC value) and therefore to determine the best temperature proxy for each hormonal response (but see for DIC criticism: Plummer, 2008; Spiegelhalter et al., 2002).

#### Population-level corticosterone responses to temperature and individual differences in corticosterone concentrations

To understand the factors that explain the population-level variation in baseline and stress-induced corticosterone, we built two univariate general linear mixed-effects models for baseline and stress-induced corticosterone, respectively, assuming a Gaussian error distribution and dividing concentrations by 1000. As fixed effects, we fitted sex (factor with two levels: male or female), year of sampling (factor with five levels: 2015-2019, to control for among-year differences), bleeding time (continuous variable, to control for the time needed to extract a blood sample, see data range above) and temperature at capture (continuous variable ranging from 3.8 to 26.8 °C). Bleeding time and temperature at capture were mean centered and variance standardized to facilitate the interpretation of their relative influence on baseline and stress-induced corticosterone. In each model, we also fitted random intercepts for individual identity to control for repeated measures of individuals across years and estimate the variance explained by among-individual differences (V_I_) and the remaining (residual, V_R_) variance. Repeatability (R) of each trait, conditional to the variance explained by the fixed effects, was estimated as the proportion of the total phenotypic variance explained by individual variance.

#### Within-individual changes in response to temperature

To analyze whether individuals changed corticosterone concentrations over their sampling period in response to ambient temperature, we used a “within-subject centering” approach (i.e., van de Pol & Verhulst, 2006; van de Pol & Wright, 2009). We calculated a) the average value of temperature that each individual has experienced and b) the observation’s deviation from the focal individual’s average value. As such, “average temperature” represents the among-individual temperature effect, while “observation’s deviation” represents the within-individual plastic change of corticosterone with temperature (van de Pol & Wright, 2009). We then built two univariate general linear mixed-effects models where either baseline or stress-induced corticosterone were fitted as response variable modelled with a Gaussian error and divided by 1000, and the among- and within-individual components of temperature were fitted as fixed effects. We modelled the same fixed and random effect structure as described above Finally, we tested whether the among- and within-individual effects of temperature at capture on corticosterone differed statistically. To do so, we calculated the difference between the parameter estimates of the within- and among-individual effect of temperature, and assessed whether their 95% Credible Intervals (CI) overlapped zero.

#### Individual differences in corticosterone plasticity

To test whether baseline and stress-induced corticosterone showed differences among individuals in their plastic response to ambient temperature at capture (individual by environment interactions, IxE), we used a random regression analysis (Nussey et al., 2007). We fitted two univariate general linear mixed-effects models with baseline and stress-induced corticosterone as response variables, respectively, assuming a Gaussian error distribution and divided by 1000. We modelled the same structure of fixed and random effects as described above, while adding the interaction of individual identity with temperature as a random slope effect. With this model, we tested for among-individual variance in the elevation (Velevation) and slopes of corticosterone concentrations in response to temperature at capture (Vslopes), while also testing for the covariance and correlation among individuals’ elevation and slopes (COVelevations-slopes and relevations-slopes, respectively). Inappropriate modelling of residual variance (e.g., assuming residual homogeneity) can lead to erroneous inferences of slope variance in random regression models (for further detail, see Ramakers et al., 2020). Hence, we assumed residual effects to be year-specific (i.e., estimated residual variance for each of the 5 study years) and uncorrelated across years (i.e., diagonal residual error structure). To assess whether models with i) both random intercept and random slope terms and/or ii) with heterogeneous residual structure explained the data better than models with only random intercepts and/or homogenous residuals, we used DIC model comparison (being the best-fitting model the one with the lowest DIC value, but see Plummer, 2008; Spiegelhalter et al., 2002 for criticism on DIC model comparisons).

#### Correlations between baseline and stress-induced corticosterone

We fitted a bivariate general linear mixed-effects model to investigate whether and how baseline and stress-induced corticosterone covaried at the population (phenotypic), among- and within-individual levels. To do so, we simultaneously modelled baseline and stress-induced corticosterone as response variables, assuming a Gaussian error distribution and divided by 1000. We fitted the same fixed and random effect structure as described above, while also fitting unstructured (co)variance matrices for the random effect individual identity and residual variance.

#### Statistical model implementation

All statistical models were fitted using a Bayesian framework implemented in the statistical software R (v. 3.6.1, R Core Team 2019) using the R-package MCMCglmm (Hadfield, 2010). For all models, we used parameter-expanded priors (Hadfield, 2010). The number of iterations and thinning interval were chosen for each model to ensure that the minimum MCMC effective sample sizes for all parameters were 1000. Burn-in was set to a minimum of 5000 iterations. The retained effective sample sizes yielded absolute autocorrelation values lower than 0.1 and satisfied convergence criteria based on the Heidelberger and Welch convergence diagnostic. We drew inferences from the posterior modes and 95% Credible Intervals (CI). We considered fixed effects and correlations to be important if the 95% CI did not include zero; estimates centered on zero were considered to provide support for the absence of an effect.

## Acknowledgments

We are indebted to Sabine Jörg and Robert de Bruijn for their field and lab help, and to Julia Cramer, Natalia Perez-Ruiz, Nataly Hidalgo Aranzamendi, Sam Hardman, Holland Galante and Skylar Buckingham for field assistance. Many thanks to the Erzdiozöse München und Freising, especially K. Meindl, B. Vollmar and M. Laußer for allowing us to work in their forest. The Hau-Goymann lab group provided input that greatly improved data collection and manuscript preparation. This work was funded by the Max-Planck-Society.

## Ethics

All experimental procedures were conducted according to the legal requirements of Germany and were approved by the governmental authorities of Oberbayern, Germany.

## Data availability

The dataset generated for this study can be found in xxx (https://…; will be supplied following acceptance for publication).

