## Supplemental Material for "Great tits differ in glucocorticoid plasticity in response to spring temperature"

SUPPLEMENTARY MATERIAL

**Table S1.** DIC model comparison of a series of univariate mixed-effects models for baseline corticosterone in response to different metrics of ambient temperature.

| Temperature variable | DIC | ΔDIC |
|---|---|---|
| Temperature at Capture | 2092.43 | 0.00 |
| Max temperature of capture day | 2097.08 | 4.65 |
| Mean temperature of capture day | 2097.37 | 4.94 |
| Min temperature of capture day | 2099.69 | 7.26 |
| Max temperature of previous day | 2101.68 | 9.25 |
| Mean temperature of previous day | 2101.72 | 9.29 |
| Max temperature on 3 previous day | 2103.03 | 10.61 |
| Min temperature of previous day | 2103.61 | 11.18 |
| Mean temperature on 3 previous day | 2103.73 | 11.31 |
| Min temperature on 3 previous day | 2105.12 | 12.70 |

**Table S2.** DIC model comparison of a series of univariate mixed-effects models for stress-induced corticosterone in response to different metrics of ambient temperature.

| Temperature variable | DIC | ΔDIC |
|---|---|---|
| Temperature at Capture | 2866.596 | 0.00 |
| Max temperature of capture day | 2873.062 | 6.47 |
| Mean temperature of capture day | 2874.68 | 8.08 |
| Max temperature of previous day | 2875.419 | 8.82 |
| Mean temperature of previous day | 2877.988 | 11.39 |
| Min temperature of capture day | 2879.085 | 12.49 |
| Max temperature on 3 previous day | 2882.943 | 16.35 |
| Mean temperature on 3 previous day | 2884.294 | 17.70 |
| Min temperature of previous day | 2884.61 | 18.01 |
| Min temperature on 3 previous day | 2886.855 | 20.26 |

**Supplementary Figure S1**

Environmental temperatures (°C, y-axis) during the breeding seasons of 2015-2019. X-axis shows Julian dates (Day 1 = Jan 1, plotted are Apr 24 – Jul 9 for each year). Symbols denote temperature averages over 7 days, with black squares and lines showing weekly daily temperatures, orange circles and lines weekly maximal temperature and blue circles and line weekly minimal temperatures.

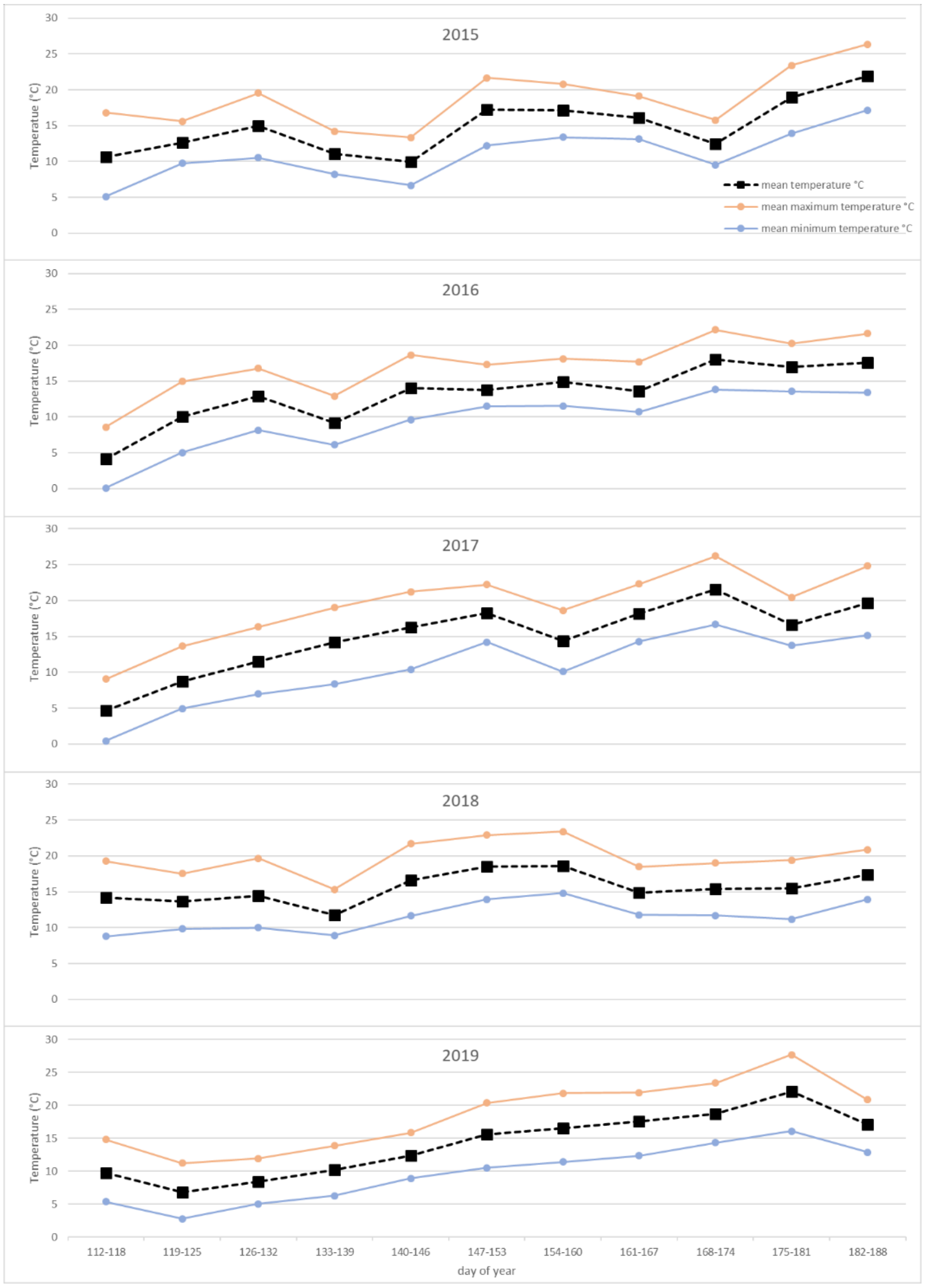
